# Human CCR4 deadenylase homolog Angel1 is a Non-Stop mRNA Decay factor

**DOI:** 10.1101/2022.04.28.489582

**Authors:** Tim Nicholson-Shaw, Megan E. Dowdle, Yasmeen Ajaj, Mark Perelis, Amit Fulzele, Gene W. Yeo, Eric J. Bennett, Jens Lykke-Andersen

**Affiliations:** Division of Biological Sciences, University of California San Diego, 9500 Gilman Drive, La Jolla, CA 92093, USA; Department of Cellular and Molecular Medicine, University of California San Diego, 9500 Gilman Drive, La Jolla, CA 92093, USA; Stem Cell Program, University of California San Diego, Sanford Consortium for Regenerative Medicine, 2880 Torrey Pines Scenic Drive, La Jolla, CA 92037, USA; Institute for Genomic Medicine, University of California San Diego, 9500 Gilman Drive, La Jolla, CA 92093, USA

## Abstract

Translation elongation stalls trigger mRNA decay and degradation of the nascent polypeptide via translation-dependent quality control pathways. One such pathway, non-stop mRNA decay (NSD), targets aberrant mRNAs that lack stop codons for example due to premature polyadenylation. Here we identify Angel1, a CCR4 deadenylase homolog whose biochemical activity remains poorly defined, as a rate-limiting factor for NSD in human cells. Angel1 associates with mRNA coding regions and proteins involved in ribosome-associated quality control and mRNA decay, consistent with a factor that monitors translation elongation stalls. Depletion of Angel1 causes stabilization of reporter mRNAs that are targeted for NSD by the absence of stop codons, but not an mRNA targeted for nonsense-mediated decay. A conserved catalytic residue of Angel1 is critical for its function in NSD. Our findings identify Angel1 as a human NSD factor and suggest that Angel1 catalytic activity plays a critical role in the NSD pathway.

## Introduction

Faithful and accurate expression of the cell’s repertoire of protein coding genes is vital to proper cellular function. However, certain cell conditions and problematic mRNA substrates can, if unresolved, lock up translational machinery in unproductive events or create potentially toxic non-functional protein products. Quality control pathways for these aberrant translation events are necessary to maintain the integrity of the proteome (Ermolaeva et al. 2015; Doma and Parker 2007; Chen et al. 2011; Lykke-Andersen and Bennett 2014; D’Orazio and Green 2021). One such pathway is activated upon the stalling of ribosomes during translation elongation and is sensed when trailing ribosomes collide with the stalled ribosome (Joazeiro 2019; Simms et al. 2017). The collided ribosomes create a unique surface that is recognized by the E3 ubiquitin ligase ZNF598 (Hel2 in the budding yeast *Saccharomyces cerevisiae*) (Juszkiewicz et al. 2018; Ikeuchi et al. 2019). This activates a cascade of events that results in degradation of the mRNA, release of the stalled ribosome, and, by a process known as Ribosome-associated Quality Control (RQC), degradation of the nascent polypeptide (D’Orazio and Green 2021).

mRNAs targeted for decay by stalled ribosomes are broadly categorized into two pathways for historical reasons rather than any obvious mechanistic distinction. One is No-Go Decay (NGD), which is defined by translational stalls that occur in the coding region (Doma and Parker 2006). NGD can be induced by strong RNA structures including G-quadruplexes (Bao et al. 2020; Endoh and Sugimoto 2016), certain amino-acid tracts (Huter et al. 2017), or oxidative damage (Simms et al. 2014). The other pathway is Non-Stop Decay (NSD) in which the ribosome stalls at the end of the mRNA because it never encounters a stop codon (Frischmeyer et al. 2002; Van Hoof et al. 2002). This can occur due to mRNA truncation (Pisareva et al. 2011) or an improper polyadenylation event, such as at a cryptic poly(A) site in the coding region. The latter leads to translation into the poly(A) tail, which causes a stall due to inefficient tRNA loading impairing translation elongation (Chandrasekaran et al. 2019). In both pathways, trailing ribosomes collide with the stalled ribosome, which triggers the subsequent degradation events.

Our best understanding of the NSD and NGD pathways comes from budding yeast. It has been long understood that decay of NGD substrates in budding yeast involves endonucleolytic cleavage at the site of the stall (Doma and Parker 2006). The endonuclease responsible for this cleavage event has been identified as Cue2 in budding yeast and its homolog NONU-1 in the worm *Caenorhabditis elegans* (D’Orazio et al. 2019; Glover et al. 2020). Endonucleolytic cleavage is also believed to be an initiating event for NSD substrates (Glover et al. 2020), but cleavage would occur close to the mRNA 3’ end which is technically difficult to monitor. The 5’ RNA fragment is subsequently degraded in a process dependent on the cytoplasmic Exosome and its associated Ski complex, whereas the 3’ fragment is degraded in a manner dependent on the 5’-to-3’ exonuclease Xrn1 (Frischmeyer et al. 2002; Tsuboi et al. 2012). In addition, NGD mRNA substrates in budding yeast have also been observed to undergo degradation independently of endonucleolytic cleavage in an Xrn1-dependent manner (D’Orazio et al. 2019).

The NSD and NGD pathways are poorly understood in mammals. Ribosome collisions or pauses appear to be common in mammals and occur at predictable motifs (Han et al. 2020). There is also evidence in human cells for large quantities of mRNAs that have undergone cleavage in a ribosome-dependent manner (Ibrahim et al. 2018). Furthermore, NSD mRNA substrates have been shown to be unstable in human HeLa cells and require the ribosome rescue factors HBS1 and DOM34, and the Exosome-SKI complex for decay (Saito et al. 2013).

A central factor in eukaryotic mRNA degradation is the CCR4-NOT deadenylase complex, which contains two catalytically active deadenylases, CCR4 (CNOT6/6L in human) and CAF1 (CNOT7/8 in human). Metazoans encode multiple homologs of CCR4, including two Angel proteins: Angel1 and Angel2 (Goldstrohm and Wickens 2008; Kurzik-Dumke and Zengerle 1996). In contrast to CCR4, no deadenylase activity was observed in biochemical assays for Angel2, which was instead found to hydrolyze 2’,3’ cyclic phosphates (Pinto et al. 2020). Angel1 was similarly found to possess 2’,3’ cyclic phosphatase activity albeit with less activity than Angel2 (Pinto et al. 2020).

In this work, we identify CCR4 deadenylase homolog Angel1 as a factor in human NSD. Angel1 associates with proteins known to be involved in NSD/NGD and RQC and with mRNA coding regions near sequences that are associated with ribosome stalling. Depletion of Angel1 causes stabilization of reporter mRNAs targeted for NSD, and mutation of a conserved catalytic residue of Angel1 disrupts this function. These observations demonstrate that Angel1 functions in human NSD and suggest that Angel1 catalytic activity is a rate-limiting step in the pathway.

## Results

### Angel1 associates with components of the RQC pathway

To gain insight into possible functions for Angel1, we first established an assay to identify Angel1 protein binding partners. Using the Flp-In T-REx system, we constructed stable human embryonic kidney (HEK) 293 cell lines expressing N-terminally FLAG-tagged Angel1 under the control of a tetracycline-regulated promoter that we titrated to express Angel1 at close to endogenous levels (Supplementary Figure S1A; see also Materials and Methods). We performed immunoprecipitation (IP) against the FLAG-tag (Supplementary Figure S1B) and identified associated proteins by liquid chromatography followed by tandem mass spectrometry (LC-MS/MS). IPs were performed with or without prior RNase A treatment to help distinguish between RNA-dependent and -independent interactions. To identify interactions specific to Angel1, the IPs were normalized to IPs from a parental Flp-In T-REx cell line expressing no FLAG-tagged fusion protein and compared to a cell line expressing FLAG-tagged TOE1, a better understood DEDD-type deadenylase with a role in snRNA processing (Lardelli et al. 2017) (Figure 1A).

**FIGURE 1.**
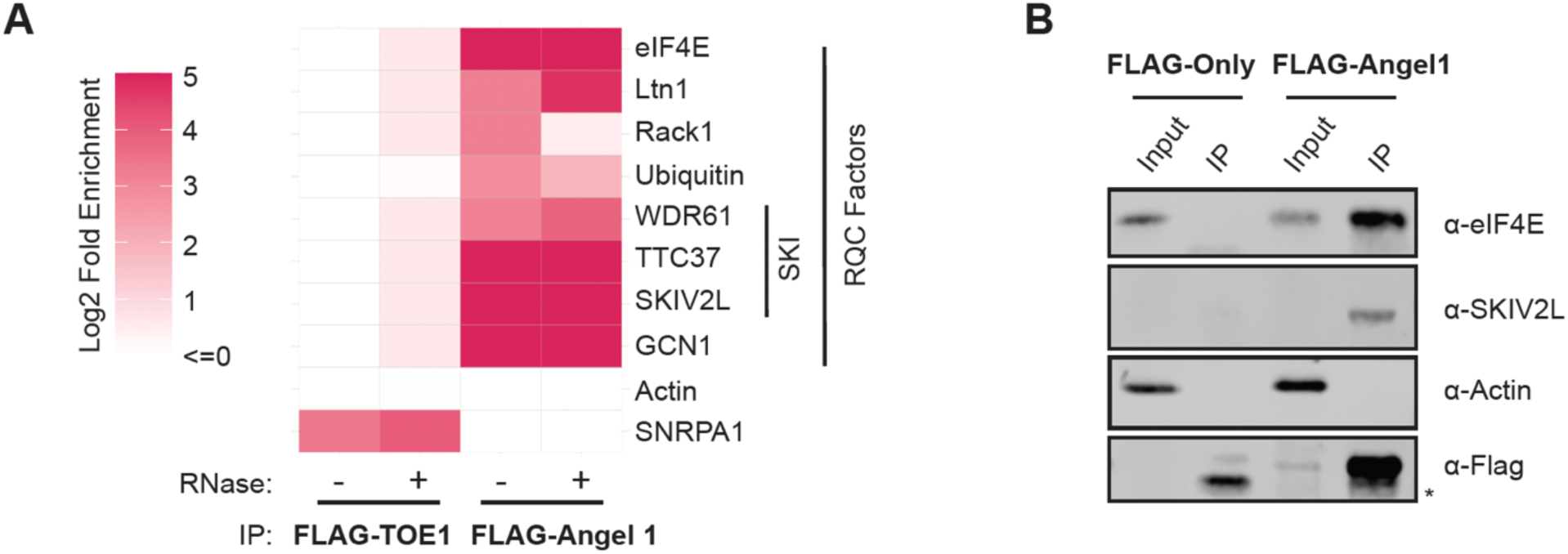
Angel1 associates with components of the NSD/NGD and RQC pathways. (A) Select proteins enriched in FLAG immune-complexes for FLAG-tagged Angel1 or TOE1 over an IP performed with a cell line expressing no FLAG-tagged protein as determined by mass spectrometry. IPs were performed in the absence (-) or presence (+) of RNase A. Fold enrichment was calculated as number of peptides per 10,000 total observed in the test IP over the negative control IP after adding a pseudocount of 1 to each identified protein. See also Supplementary Table S1. (B) Co-IP assays followed by Western blotting monitoring specific proteins associated with Angel1. Actin served as a negative control. Input: 10% of the total cell extract used for IP. *: non-specific band.

Among the most abundant proteins that specifically co-purified with FLAG-tagged Angel1 (Supplementary Table S1), was the mRNA cap-binding protein eIF4E, which, importantly, reproduces a previously described interaction (Gosselin et al. 2013). Other proteins that specifically co-purified with Angel1 included additional mRNP components (LARP4, LARP4B, DDX6, LSM14A, ATXN2, and PABPC), all components of the GATOR2 complex (MIOS, WDR24, WDR59, and SEH1L) involved in activation of mTORC1 (Cai et al. 2016), and components of a complex important for cytoskeletal functions of neurons and synaptic plasticity (DISC1, NDE1 and NDEL1; (Tropea et al. 2018).

A striking subset of Angel1-associated proteins were components of the NSD/NGD and RQC pathways (Figure 1A). These included RACK1, LTN1, GCN1, ubiquitin, and all three components of the SKI2-3-8 complex (known as SKIV2L, TTC37 and WDR61 in human), an RNA helicase complex associated with the cytoplasmic RNA Exosome. With the exception of RACK1, these all associated with Angel1 in a manner resistant to RNase A treatment (Figure 1A), suggesting protein-mediated interactions that are independent of RNA. We confirmed the association of Angel1 with eIF4E and the SKIV2L subunit of the SKI complex by IP followed by immunoblotting (Figure 1B). Given the homology of Angel1 with 3’ RNA processing factors and its association with components of the NSD/NGD and RQC pathways, we explored the hypothesis that Angel1 is involved in quality control-dependent degradation of mRNAs associated with stalled ribosomes.

### Angel1 associates with mRNA coding regions and sequence features correlated with stalled ribosomes

We were interested in understanding what RNA transcripts and sequence motifs Angel1 interacts with, reasoning that Angel1 may show preference for regions of transcripts associated with stalled ribosomes. To that end, we performed enhanced cross-linking and immunoprecipitation followed by sequencing (eCLIP-seq) (Van Nostrand et al. 2016). Two replicates of FLAG-Angel1 eCLIP-seq were performed on our Flp-In T-REx HEK293 cell lines expressing Angel1 at close to endogenous levels. We also performed eCLIP-seq with the parental cell line as a background control. When evaluating differential enrichment of genes in IPs over input, we found high agreement between the two Angel1 eCLIP replicates (Figure 2A and Supplementary Figures 2A and 2B). Angel1 eCLIP reads were significantly enriched for coding regions of mRNAs and depleted for intronic regions (Figures 2B and 2C) which is consistent with a factor that may monitor translation. Angel1 eCLIP also showed an enrichment for reads mapping to 5’UTRs (Figure 2C), which is another region of mRNAs engaged with ribosomal subunits and consistent with the association of Angel1 with eIF4E. Using an eCLIP-Seq analysis pipeline (Van Nostrand et al. 2016), we identified CLIP peaks for each replicate (p<0.05). Limiting the analysis to CLIP peaks that were reproducible between the two replicates (cutoff threshold p < 0.001) showed a further increase in the percentage of peaks mapping to coding regions (Figure 2B).

**FIGURE 2.**
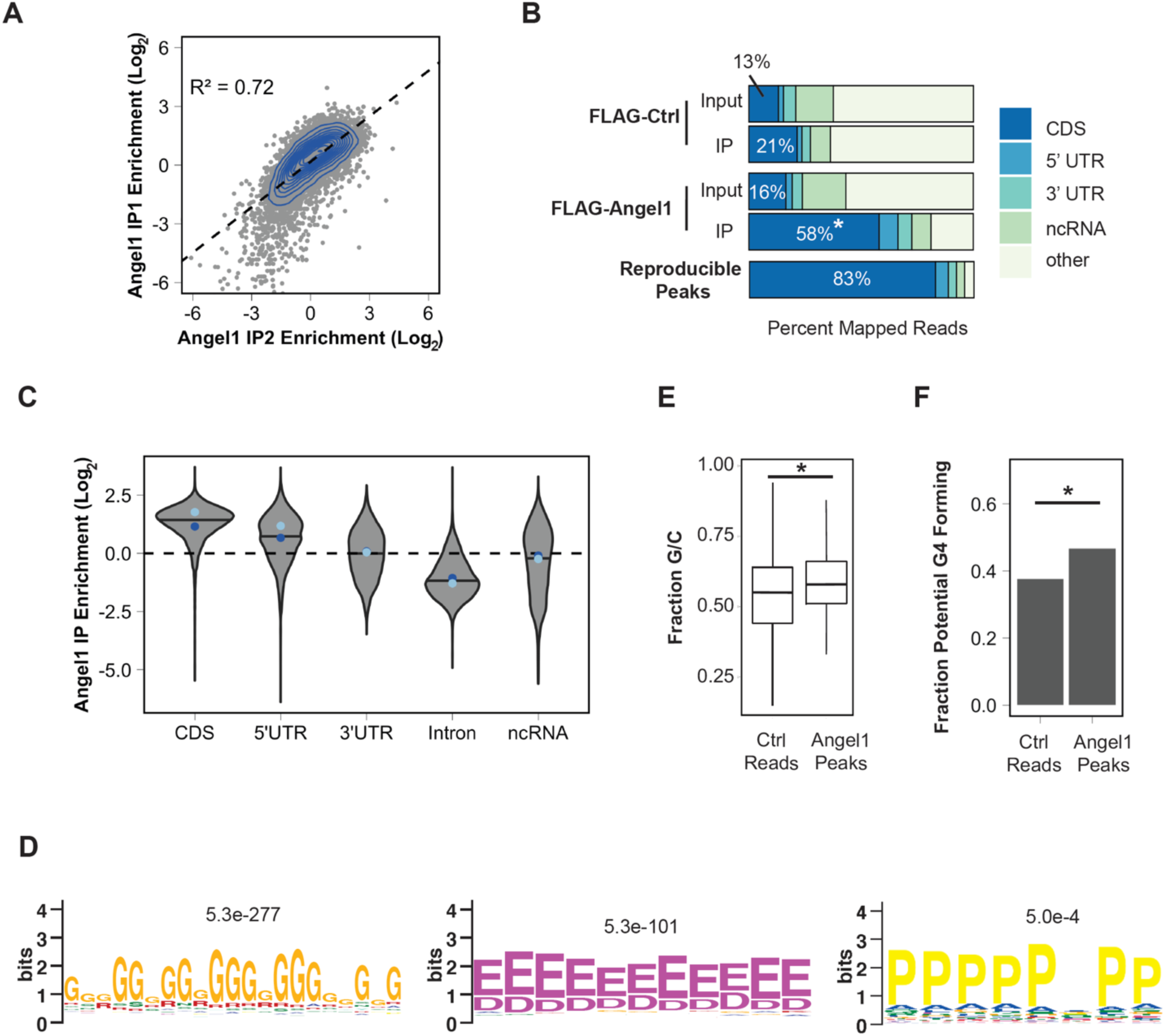
Angel1 associates with coding regions of mRNAs and with sequences associated with ribosomal stalling. (A) Log_2_ fold enrichment of IP reads over input compared between the two Flag-tagged Angel1 eCLIP replicates. Grey dots represent individual genes and the blue contour lines represent a 2D kernel density estimate. A pearson correlation is shown. (B) Fraction of reads mapping to different functional regions of RNAs in control versus FLAG-tagged Angel1 input and eCLIP (IP) samples, and in reproducible peaks (p<0.001) between the two FLAG-tagged Angel1 eCLIP experiments. (C) Fold enrichment violin plot distributions separated by annotation. The line within each distribution represents the median averaged between each replicate and the median of each individual replicate is shown with a dot. IP1: light blue, IP2: dark blue. (D) Peptide motifs enriched in areas within 50 nucleotides upstream or downstream of identified peaks as compared to areas around reads from the control sample. E-values are listed. The two leftmost shown motifs were the most highly enriched motifs in the MEME analysis. (E) GC content of sequences between 50 nucleotides upstream or downstream of identified Angel1 eCLIP peaks as compared to areas around reads from the control sample. *: p<2.2e-22 (KS-test). (F) Calculated guanosine quadruplex (G4) formation capacity of sequences between 50 nucleotides upstream or downstream of identified Angel1 eCLIP peaks as compared to areas around reads from the control sample. *: p<2.2e-22 (KS-test).

Typically, nucleotide motifs for RNA binding proteins are identified by applying motif finding algorithms to the sequences of reproducible peaks. However, this analysis failed to produce strong nucleotide or codon motifs in Angel1-associated peaks. We reasoned that, if involved in translation elongation quality control, Angel1 might be recruited to regions upstream or downstream of ribosome stalls and therefore examined regions within 50 nucleotides upstream and downstream of each peak. Motif analysis (Bailey et al. 2009) of these Angel1-associated regions revealed an abundance of guanosine-rich sequences (Supplementary Figure S2C) and several amino acid-coding motifs that have been associated with stalled ribosomes (Chyzyńska et al. 2021), including poly-glycine and poly-glutamate/aspartate—which were the two most significantly enriched motifs in the analysis—and, less enriched, poly-proline codons (Figure 2D). We also examined nucleotide content of regions around peaks and found that they contained, on average, higher GC content than regions surrounding reads in the control samples (Figure 2E) and were more likely to contain what are predicted to be more stable RNA structures (Supplementary Figure S2D). Using a G-quadruplex prediction algorithm (Kikin et al. 2006), we found that regions around peaks were also predicted to more likely form G-quadruplexes (Figure 2F), a secondary structure element that has also been associated with ribosome stalling (Endoh and Sugimoto 2016). These associations are consistent with a factor recruited to the wide variety of nucleotide and nascent oligopeptide sequences that may induce ribosome stalling.

### Development of a human NSD assay

While several factors involved in mRNA degradation by NSD and NGD have been identified in budding yeast (D’Orazio et al. 2019; Van Hoof et al. 2002; Frischmeyer et al. 2002; Tsuboi et al. 2012) and in *C. elegans* (Glover et al. 2020), much less is known about factors in human cells (Saito et al. 2013). To establish an assay to monitor the NSD pathway in human cells, we adapted the well-characterized β-globin mRNA pulse-chase system (Lykke-Andersen et al. 2000) by generating an NSD reporter mRNA lacking stop codons. In this system, wild-type β-globin mRNA is highly stable with a half-life of over 600 minutes (Durand and Lykke-Andersen 2013). Removal of all stop codons created an unstable mRNA (BG-NSD) that is degraded at a rate faster than the well-characterized β-globin Nonsense-Mediated Decay (NMD) reporter mRNA containing a premature termination codon at position 39 (BG-NMD) (Durand et al. 2016) (Figures 3A and 3B). Importantly, a single point mutation that reintroduces a stop codon nine nucleotides upstream of the cleavage and polyadenylation site (BG-NSD+Stop) stabilized the BG-NSD reporter (Figures 3A and 3B). These substrates provide a platform for investigating the effects of perturbations of human NSD machinery on mRNA decay rates.

**FIGURE 3.**
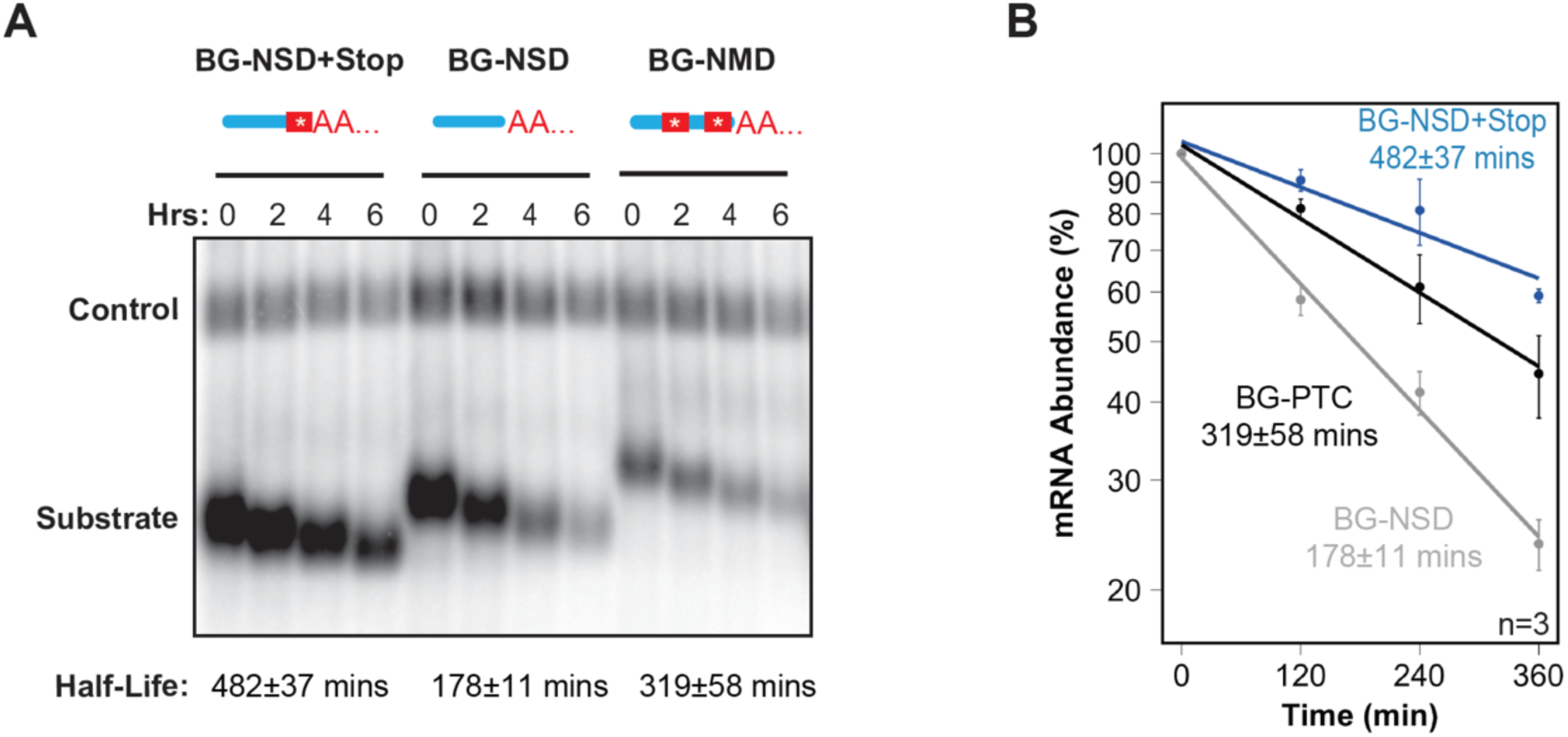
Establishment of a human NSD reporter mRNA decay assay. (A) Representative Northern blot of a pulse-chase mRNA decay assay in HeLa Tet-off cells monitoring degradation of BG-NSD+STOP, BG-NSD and BG-NMD mRNAs (Substrate) as compared with constitutively expressed β-globin-GAP3 control mRNA (Control). Numbers above lanes refer to hours after transcription shut-off of the substrate mRNAs by tetracycline. Bands were quantified and normalized to the constitutively expressed β-globin-GAP3 mRNA to calculate mRNA half-lives assuming first-order kinetics, which are given below the blot with standard error of the mean from three experiments. (B) Exponential decay plots of the experiment in panel *A* performed in triplicate (n=3). Error bars represent standard error of the mean.

To further validate our NSD reporter mRNA we tested the effect of depleting SKIV2L, a component of the human SKI complex with an established function in NSD (Saito et al. 2013). As expected, SKIV2L depletion led to stabilization of the NSD reporter (Figure 4A and Supplementary Figures S3A and S3B). We next tested whether N4BP2, a mammalian homolog of the initiating NSD/NGD endonuclease Cue2/NONU-1, plays a role in targeting the NSD substrate for mRNA decay. Indeed, depletion of N4BP2 led to stabilization of the NSD reporter (Figure 4B and supplementary Figure S3A). In contrast, depletion of XRN1, using knock-down conditions that stabilize a cleavage intermediate in the NMD pathway (Franks et al. 2010) (Supplementary Figure S3B), did not stabilize the β-globin NSD substrate (Figure 4C), likely reflecting that our assay is unable to monitor the fate of the portion of the RNA located downstream of the ribosome stalling site. These observations identify SKIV2L and N4BP2 as rate-limiting for the degradation of the NSD reporter mRNA, and suggest that N4BP2 is the human ortholog of the *S. cerevisiae* and *C. elegans* NSD/NGD endonuclease.

**FIGURE 4.**
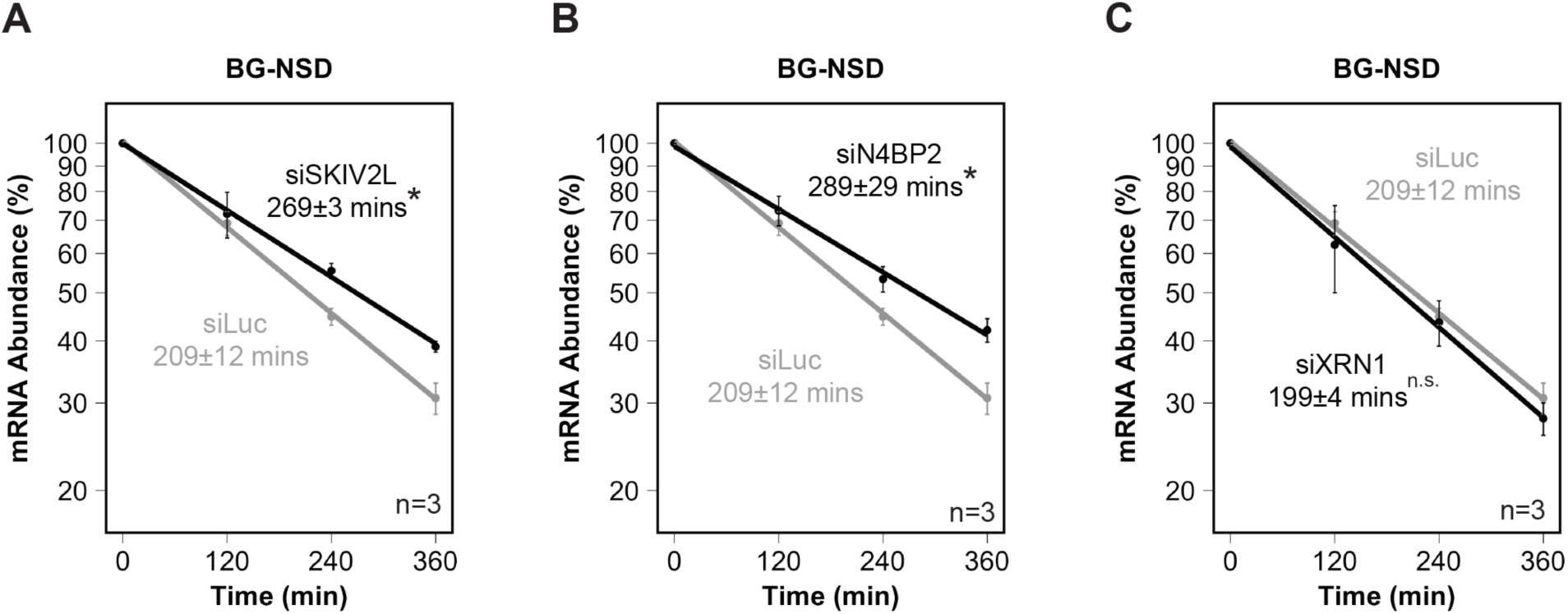
Depletion of SKIV2L and N4BP2 stabilizes the human NSD reporter mRNA. (A) Exponential decay plots of the BG-NSD substrate after depletion of known NSD factor SKIV2L. (B) Exponential decay plots of the BG-NSD substrate after depletion of N4BP2. (C) Exponential decay plots of the BG-NSD substrate after depletion of XRN1. Error bars represent standard error of the mean (n=3). *: p<0.05, calculated by one-tailed Student’s t-test compared to the control knockdown targeting luciferase (siLuc).

### Angel1 is limiting for NSD

We next tested whether Angel1 contributes to human NSD. Indeed, siRNA-mediated depletion of Angel1 (Supplementary Figure S4A) resulted in stabilization of the β-globin NSD reporter mRNA (BG-NSD) (Figure 5A). This effect was observed for two independent siRNAs targeting Angel1 (Supplementary Figure S4B). By contrast, depletion of Angel1 did not alter the stability of the β-globin NMD reporter mRNA (BG-NMD) (Figure 5B), showing that the effect is specific to turnover of the NSD substrate and not due to general repression of translation-dependent mRNA turnover. Depletion of Angel1 also resulted in stabilization of an NSD reporter based on TPI mRNA (Supplementary Figure S4C), showing that the effect of Angel1 is not specific to a β-globin mRNA substrate.

**FIGURE 5.**
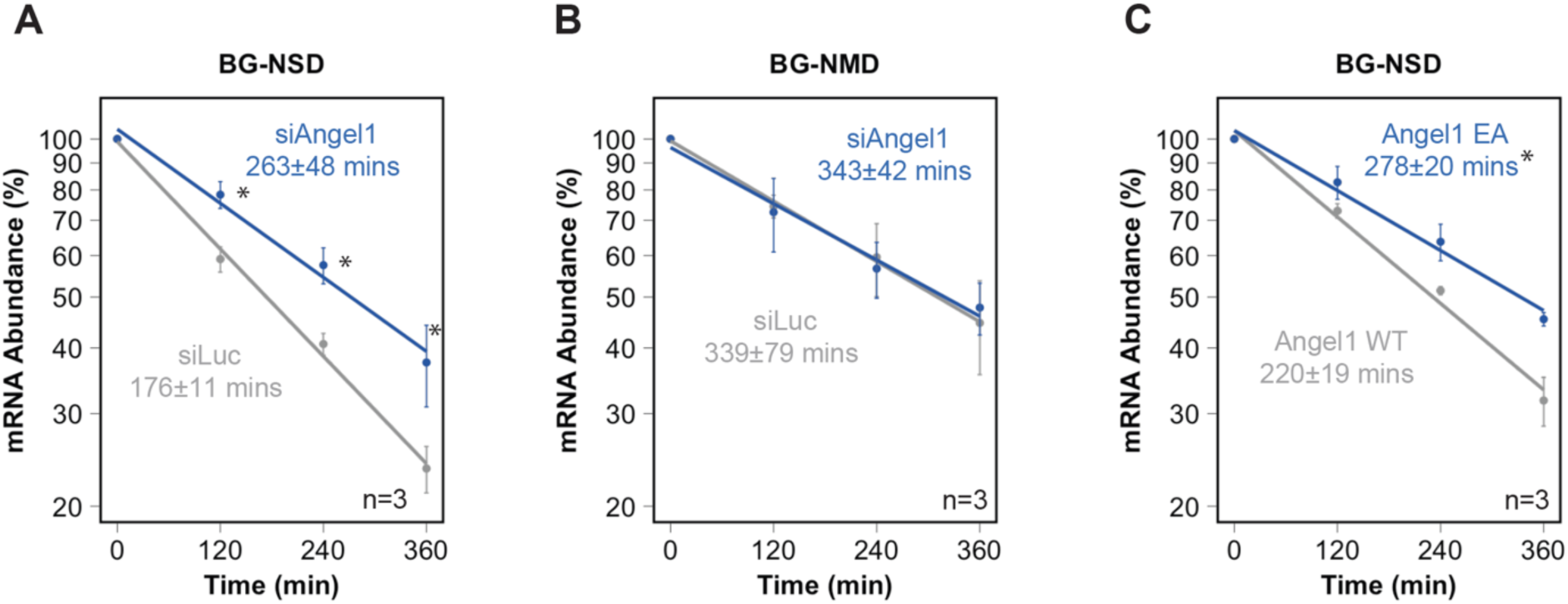
Angel1 and its catalytic center is rate-limiting for the degradation of an NSD target mRNA. (A) Exponential decay plots of the BG-NSD substrate after depletion of Angel1 or using a control siRNA (siLuc). (B) Exponential decay plots of a substrate containing a premature termination codon that is targeted for Nonsense-Mediated Decay (BG-NMD) after depletion of Angel1 or using a control siRNA. (C) Exponential decay plots of the BG-NSD substrate after depletion of Angel1 and complementing with exogenous Angel1 WT or catalytic site mutant Angel1 EA. *: p<0.05 calculated by a one-tailed Student’s t-test. Error bars represent standard error of the mean (n=3).

While the biochemical function of Angel1 remains poorly defined, catalytic residues found in the homologous CCR4 deadenylases and in the Angel2 2’,3’ cyclic phosphatase are conserved in Angel1. To test whether the conserved Angel1 catalytic center is important for its function in NSD, we generated an Angel1 protein containing a glutamate to alanine substitution previously shown to disrupt 2’,3’ cyclic phosphatase activity of the Angel1 homolog Angel2 (Pinto et al. 2020) and deadenylase activity of CCR4 (Wang et al. 2010; Yi et al. 2018). Exogenous expression of wild-type Angel1 partially rescued the effect of Angel1 depletion on NSD reporter stability (Figure 5C and Supplementary Figure S4D). By contrast, no rescue of activity was observed upon expression of the catalytic site mutant Angel1 (Angel1 EA) despite expression at equal levels as wild-type Angel1 (Supplementary Figure S4D), demonstrating that the activity of Angel1 in NSD depends on a central catalytic residue.

## Discussion

The mechanism by which mRNAs subjected to NSD and NGD are degraded in human cells remains poorly defined. In this work, we present evidence that CCR4 deadenylase homolog Angel1 facilitates decay of human mRNAs targeted for NSD. Indeed, Angel1 associates with proteins involved in the related NSD/NGD and RQC pathways (Figure 1) and with coding regions of mRNAs, including RNA sequences that have been associated with translational stalling (Figure 2). Depletion of Angel1 stabilizes NSD reporter mRNAs, and a conserved catalytic residue is critical for this activity (Figures 3-5). Thus, Angel1 is a human NSD factor.

### By what mechanism does Angel1 facilitate NSD?

Angel1 and Angel2 are a homologs of EEP-type CCR4 deadenylases, but we (Supplementary Figure S5) and others (Pinto et al. 2020) have observed no evidence for deadenylase activity of human Angel proteins in biochemical assays. Instead, Angel2, and to a lesser degree Angel1, was found to have activity as a 2’,3’ cyclic phosphatase, dependent on a highly conserved catalytic glutamate residue (Pinto et al. 2020). Our observation that Angel1’s activity in NSD is dependent on this conserved catalytic glutamate residue suggests that Angel1 has a catalytic function in the NSD pathway, although we cannot formally rule out a structural defect caused by the mutation. The endonucleolytic cleavage event in NSD/NGD that is catalyzed by Cue2/NONU-1/N4BP2 is predicted to generate a 2’,3’ cyclic phosphate at the 3’ end of the 5’ RNA fragment (Yang 2011; Navickas et al. 2020). This cyclic phosphate is a potential catalytic target of Angel1, which could help facilitate subsequent degradation of the 5’ fragment. However, we observed no detectable accumulation of a 2’,3’ cyclic phosphate in our BG-NSD reporter mRNA upon depletion of Angel1 in an assay capable of detecting a 2’,3’ cyclic phosphate generated by a ribozyme in the same reporter mRNA (Supplementary Figure S6). Another potential substrate for a 2’,3’ cyclic phosphatase in the NSD pathway could be the P-site tRNA which is left with a 2’,3’ cyclic phosphate after removal and degradation of the nascent polypeptide (Yip et al. 2019). However, the resolution of this cyclic phosphate seems a less likely candidate to explain the observed impact of Angel1 on mRNA degradation. While a cyclic phosphatase function may be the most parsimonious explanation for the activity of Angel1 in NSD, it cannot be ruled out that Angel1 functions by a different mechanism, such as by acting as a deadenylase or 3’-to-5’ exonuclease despite the absence of such an activity in biochemical assays. Angel1 could also impact NSD by non-catalytic mechanisms such as via its interaction with factors that impact translation and mRNA stability, including eIF4E, PABPC, LSM14A and DDX6 proteins (Figure 1 and Supplementary Table S1).

### Possible functions for Angel1 outside of RQC

In addition to RQC factors, we also found association in our IP-MS/MS analysis of Angel1 with the Gator2 complex which, along with Sestrins and Gator1, is important for sensing amino acid deprivation and signaling through mTORC1 (Bar-Peled et al. 2013; Kowalsky et al. 2020). The association of Angel1 with these components suggests a potential role in sensing or modulating amino acid deprivation. Given Angel1’s homology to 2’,3’ cyclic phosphatase Angel2, such a function could be related to tRNAs which can be cleaved during tRNA splicing and stress conditions to create 2’,3’ cyclic phosphate-containing species (Zillmann et al. 1991; Shigematsu and Kirino 2020). Furthermore, Angel1’s association with DISC1-NDE1/NDEL1 implicates Angel1 in cytoskeletal functions, although these proteins have no currently known role in RNA metabolism. Angel1 could also be involved in additional processes that involve cyclic phosphates, such as the metabolism of RNAs that feature cyclic phosphates during their life-cycles, including U6 snRNA (Gu et al. 1997), spliced tRNAs (Zillmann et al. 1991), or XBP1 mRNA (Jurkin et al. 2014).

### What are the endogenous substrates of Angel1 and the human NSD/NGD pathway?

While our reporter assays show that Angel1 is rate-limiting for decay of engineered human NSD substrates (Figure 5), our eCLIP experiments suggest that Angel1 associates broadly with protein coding regions of mRNAs (Figure 2). Indeed, depletion of RQC factors such as ZNF598 have shown broad, low-level, effects on the transcriptome (Tuck et al. 2020; Kalisiak et al. 2017; Weber et al. 2020; Sundaramoorthy et al. 2021). Identification of endogenous substrates of RQC has been generally unsuccessful with only a few potential examples, including the ER stress-induced XBP1, which in human cells is upregulated at the protein level upon depletion of ZNF598 (Han et al. 2020). These observations suggest broad pleiotropic effects of perturbations in this system, perhaps reflecting a process that occurs stochastically at individual translated mRNAs under normal conditions. We also observed association of Angel1 with mRNA 5’UTRs, which is consistent with the association of Angel1 with eIF4E and could potentially relate to mRNA upstream open reading frames, or to the recently described RQC pathway detecting collisions between scanning pre-initiation complexes (Garshott et al. 2021). Altogether, our study identifies Angel1 as a factor involved in human NSD. An important question for future study is how Angel1 integrates with other NSD/NGD factors, including the endonuclease N4BP2 and the SKI2-3-8-exosome complex, to degrade NSD/NGD substrates.

## Acknowledgements

We would like to thank the members of the Lykke-Andersen lab for their input and useful discussions. We also would like to thank the Triton Shared Compute Cluster (TSCC) at the San Diego Supercomputer Center for use of their hardware for alignments. Sequencing was conducted at the IGM Genomics Center, University of California, San Diego, La Jolla, CA. This work was supported by National Institutes of Health (NIH) grant R35 GM118069 awarded to J.L-A, NIH NSRA F32 fellowship GM151845 awarded to M.E.D., NIH NRSA F32 fellowship 5F32HL143978-02 to M.P., and NIH grant R35 GM148339 awarded to E.J.B. G.W.Y. is supported by National Institutes of Health grants HG004659 and HG009889. G.W.Y. is supported by an Allen Distinguished Investigator Award, a Paul G. Allen Frontiers Group advised grant of the Paul G. Allen Family Foundation.

## Materials and Methods

### Antibodies

Western blotting was performed with anti-FLAG (Sigma F7425; 1:1,000), anti-eIF4E (Cell Signaling Technologies 9742; 1:1,000), anti-SKIV2L (Thermo Fisher 11462-1-AP; 1:500), anti-β-actin (Cell Signaling Technologies 4967; 1:1,000).

### Plasmids

Gibson assembly (New England Biolabs) was used to insert the coding region of Angel1 with an engineered N-terminal FLAG-tag into pcDNA5/FRT/TO and pcDNA3 to create pcDNA5/FRT/TO-FLAG-Angel1 WT and pcDNA3-FLAG-Angel1 WT. Site-directed mutagenesis (New England Biolabs, E0554S) was used to create an E298A catalytically inactivating mutation, generating pcDNA3-FLAG-Angel1 EA. Pulse-chase constructs were created from the previously described plasmid, pcTet2-BWT (Damgaard and Lykke-Andersen 2011). pcTet2-BG-NSD was created by 3 rounds of site-directed mutagenesis that removed all in-frame stop codons before the cleavage and polyadenylation site through deletion of a 70 bp region and 6 point mutations. A point mutation was introduced 10 nucleotides upstream of the cleavage and polyadenylation site to create a stop codon, generating pcTet2-BG-NSD+stop. Additionally, a self-cleaving ribozyme sequence from the herpes deltavirus was inserted into pcTet2-BWT generating pcTet2-BG-HDV. pcTet2-BG-NMD was created by site-directed mutagenesis introducing a stop codon at codon 39 of pcTet2-BWT. pcBGAP3 was used as an internal control for pulse-chase experiments (Clement and Lykke-Andersen 2008). pcTet2-TPI-NSD was generated from pcTet2-TPI (Singh et al. 2008) using a synthesized double stranded gene fragment (IDT, gBlock) of the 3’UTR of pcTet2-TPI edited to remove all in-frame stop codons. The gene fragment was used to replace the 3’UTR of pcTet2-TPI by Gibson assembly. Plasmid sequences are available upon request.

### Stable cell line construction and titration of FLAG-Angel1 levels

pcDNA5/FRT/TO-FLAG-Angel1 WT was used to generate stable HEK FLp-In T-REx-293 cell lines (Invitrogen) according to the manufacturer’s protocol, in which FLAG-Angel1 expression can be titrated with tetracycline. In the absence of an Angel1 antibody, we used an anti-FLAG antibody to estimate FLAG-Angel1 expression levels in comparison to FLAG-TOE1, which had been titrated to endogenous levels as monitored by a TOE1 antibody (Wagner et al. 2007; Lardelli et al. 2017). TOE1 is approximately 25 times more abundant than Angel1 in HeLa cells according to global mass spectrometry measurements (Nagaraj et al. 2011). We therefore titrated FLAG-Angel1 expression with tetracycline to match a level of approximately 1:25 relative to TOE1, which was reached at 5 ng/ml of tetracycline. This concentration of tetracycline was used in all experiments when expressing FLAG-Angel1 in the stable HEK Flp-In T-REx-293 line.

### Cell growth and depletions

Cells were maintained in Dulbecco’s Modified Eagle Medium (DMEM, Gibco, 11965092) with 10% fetal bovine serum (Gibco, 10437028). Flp-In T-REx lines were induced with 5 ng/ml tetracycline 24 hours before harvest. Knockdowns were performed using 20 nM of small interfering RNAs (siRNAs) custom ordered from Horizon Discovery (Supplementary Table S2). The control siRNA targeted luciferase mRNA. Knockdowns were performed with siLentFect reagent (Bio-Rad, 703362) according to the manufacturer’s specifications.

### Pulse-chase mRNA decay assays

Hela Tet-off cells were plated at 15×10^4^ cells per well in a 6-well plate. siRNA-mediated knockdowns were performed at 72 hours and 24 hours prior to cell harvest. In addition, 48 hours prior to cell harvest, cells were transfected with 0.5 µg of the test construct (pcTet2-BG-NSD, -BG-NSD+Stop, or -BG-NMD), 0.5 µg of pcDNA3-based Angel1 addback construct (if applicable), 0.1 µg of a pcBGAP3 loading control plasmid, and empty pcDNA3 plasmid stuffer to a total of 2 µg. Cells were maintained with 50 ng/ml tetracycline to prevent expression from the test plasmid. 72 hours after the initial siRNA transfection, cells were rinsed with PBS, and transcription from the test plasmids was pulsed by addition of 2 ml of fresh medium free of tetracycline for 6 hours. Medium was subsequently replaced with DMEM/10% FBS containing 1,000 ng/ml tetracycline to shut off test plasmid transcription and cells were collected every 2 hours thereafter in Trizol reagent (Thermo Fisher, 15596026). RNA was isolated and substrate levels were analyzed by Northern blotting as previously described (Clement and Lykke-Andersen 2008).

### Immunoprecipitation assays

Flp-In T-REx lines expressing FLAG-tagged Angel1, FLAG-tagged TOE1, or no FLAG-tagged fusion protein were grown to approximately 50% confluency and induced with 5 ng/ml tetracycline for 24 hours. Cells were harvested by scraping into ice cold PBS and flash frozen in liquid nitrogen. Pellets were resuspended in isotonic lysis buffer (50 mM Tris-HCl pH 7.5, 150 mM NaCl, 0.2 mM EDTA, 0.5% Triton X-100) with 80 units/ml RNaseOUT (Thermo Fisher, 10777019) or 125 µg/ml RNase A (Sigma, R4875), and 1 tablet/10 ml of protease inhibitor (Thermo Fisher, 88666)) for 10 minutes on ice. Lysates were spun down at 20,000 *g* for 15 minutes at 4°C. FLAG peptide (ApexBio, A6002) was added to lysates to a concentration of 1 µg/ml to reduce non-specific interactions. Samples were incubated with pre-washed anti-FLAG M2 agarose beads (Sigma, A2220) for 2 hours at 4°C with rotation. Beads were washed 9 times with NET2 buffer (10 mM Tris-HCl pH 7.5, 150 mM NaCl, 0.1% Triton X-100). Protein was eluted by treating beads three times for 30 minutes at 4°C with NET2 containing 200 µg/ml FLAG peptide and elutions were subsequently pooled. Samples from input, the unbound fraction, and elution were separated by gel electrophoresis and visualized by silver staining (Thermo Fisher, 24580) according to the manufacturer’s protocol. Protein amounts for deadenylation assays were estimated against BSA standards (New England Biolabs, B9000S).

### Mass Spectrometry analysis

In brief, the Immunoprecipitated samples were in-solution digested overnight at 37 °C in 400 ng of mass spectrometry grade trypsin (Promega) enzyme. The digestion was stopped by adding formic acid to the 0.5% final concentration. The digested peptides were desalted by using C18 StageTips and were transferred to a fresh tubes and then vacuum dried. The vacuum-dried peptides were resuspended in 5% formic acid/5% acetonitrile buffer and added to the vials for mass spectrometry analysis. Samples were analyzed with duplicate injections by LC-MS-MS using EASY-nLC 1000 liquid chromatography connected with Q-Exactive mass spectrometer (Thermo Scientific) as described previously (Sundaramoorthy et al. 2017) with some modification as follows. The peptides were eluted using the 60 min acetonitrile gradient (45 min 2%−30% ACN gradient followed by 5 min 30%− 60% ACN gradient, a 2 min 60−95% ACN gradient, and a final 8 min isocratic column equilibration step at 0% ACN) at a 250 nl/min flow rate. All the gradient mobile phases contained 0.1% formic acid. The data dependent analysis (DDA) was done using the top 10 method with a positive polarity, scan range of 400−1800 m/z, a 70,000 resolution, and an AGC target of 3e6. A dynamic exclusion time of 20 s was implemented and unassigned; singly charged and charge states above 6 were excluded for the data dependent MS/MS scans. The MS2 scans were triggered with a minimum AGC target threshold of 1e5 and with a maximum injection time of 60 ms. The peptides were fragmented using a normalized collision energy (NCE) setting of 25. Apex trigger and peptide match settings were disabled. RAW files were processed, searched, and analyzed essentially as described previously (Reinke et al. 2017). To calculate the fold enrichment of individual proteins in the Angel1 IP over the matched FLAG control, the number of peptides for each protein were normalized to counts per 10,000 in the total count for each sample, and the normalized counts for Angel1 IP were divided by normalized counts for the control after adding a pseudocount of 1 to every normalized peptide count to prevent division by zero errors. All experiment related RAW mass-spectrometry data files were deposited at the MassIVE repository using the accession identifier MassIVE: MSV000089129.

### eCLIP assays

Flp-In TREx lines expressing FLAG-tagged Angel1 or no FLAG-tagged fusion protein were grown to approximately 50% confluency and induced with 5 ng/ml tetracycline for 24 hours. Cells were crosslinked to preserve protein-RNA interactions by treatment with UV (Stratalinker, 254 nm, 400 mJ/cm^2^, on ice). One sample was not exposed to UV as a no-crosslink control. eCLIP library preparation was performed as previously detailed (Van Nostrand et al. 2017). Samples were mapped to the hg37 human genome and features from the Gencode 19 annotation were counted with featureCounts (Liao et al. 2014). Reads were annotated with a Yeo lab annotation pipeline (Annotator v0.0.13). Fold enrichment was calculated as enrichment of IP reads over input reads for genes with mapped reads in the input condition above a threshold. Areas 50 nucleotides upstream and downstream from peaks were extracted with custom python scripts that used transcripts tagged as Appris principal (Rodriguez et al. 2013) to limit genes to one transcript. In cases where genes had multiple principal transcripts, the longest transcript was selected. G/C content was calculated with custom scripts for those regions. G quadruplex formation potential was measured for those sequences using QGRS (Kikin et al. 2006). Significance was tested with a Kolmogorov-Smirnov (KS) test. Sequencing data have been deposited into the Gene Expression Omnibus (GEO) under accession number GSE199650.

### RT-qPCR assays

After total RNA isolation from cells, reverse transcription was performed using Superscript III (Thermo Fisher, 18080044) according to manufacturer’s protocol. qPCR was performed with a master mix (Thermo Fisher, 4385612) and using a Quantstudio 3 machine according to manufacturer specifications. Angel1 qPCR was carried out using pre-validated primers (Bio-Rad, 10025636). All other qPCR primer pairs are listed in Supplementary Table S2.

### Deadenylation assay

A custom poly-A RNA substrate terminating in 20 adenosines (Dharmacon) and a DNA loading control also terminating in 20 adenosines (IDT) (Supplementary Table S2) were 5’ labelled with [γ-^32^P]-ATP (PerkinElmer) using T4 poly-nucleotide kinase (NEB) according to manufacturer’s protocol. The deadenylation assay was adapted from a previously described protocol (Wagner et al. 2007). Approximately 50 nM of the indicated protein was added to approximately 5,000 CPM each of DNA loading control and RNA substrate and incubated at 37°C in deadenylation buffer (20mM HEPES, pH7.4, 2mM MgCl_2_, 0.1 mg/ml bovine serum albumin, 1 mM spermidine, 0.1% Igepal CA-630 (Sigma), 0.5 units/µl RNase-Out, and 0.5 μg/µl yeast total RNA). Formamide loading buffer was added to stop the reaction and samples were separated in a 9% polyacrylamide-6M urea denaturing gel. The gel was dried and imaged using a phosphorimager.

### 2’,3’-cyclic phosphate assay

Hela Tet-off cells were plated at 15×10^4^ cells per well in a 6-well plate. siRNA-mediated knockdowns targeting Luciferase and Angel1 were performed at 72 hours and 24 hours prior to cell harvest. In addition, 48 hours prior to cell harvest, cells were transfected with 0.5 µg of the test plasmid (pcTet2-BG-NSD, -BG-NSD+Stop, or -BG-HDV) and 1.5 µg of internal control plasmid pcPC-GPx1, which expresses rat GPx1 mRNA (Singh et al. 2007). Cells, which were maintained in the absence of tetracycline to allow β-globin mRNA expression, were then harvested with Trizol reagent and total RNA was isolated. After RNA isolation, 10 µg of RNA was DNase treated with Turbo DNase (Thermo Fisher, AM2238) according to manufacturer’s protocol. After DNase treatment, the RNA was split into two pools for 3’-end RNA adapter ligations with either RtcB ligase (NEB, M0458S) or T4 RNA ligase (NEB, M0204S). RtcB reactions were carried out to a final reaction concentration of 5 µg of DNase-treated RNA, 2 µM of Ag10-5’OH RNA adapter (Supplementary Table S2), 1x of RtcB reaction buffer, 1 mM MnCl_2_, 0.1 mM GTP, 40 units of RNaseOUT (Thermo Fisher, 10777019), 15% PEG8000, and 1 µl of 15 µM RtcB ligase, in a 20 µl total reaction volume. RtcB ligation reactions were then incubated at 37°C for 2 hours. T4 RNA Ligase reactions were carried out with final reaction concentrations of 5 µg RNA, 2 µM Ag10-5’P RNA adapter (Supplementary Table S2), 1 mM ATP, 0.04 mg BSA, 1x T4 RNA Ligase Reaction Buffer, 20 units RNaseOut, and 30 units T4 RNA Ligase 1 (NEB) in a 10 µl total reaction. T4 RNA ligation reactions were then incubated at 16°C overnight. Ligation reactions then underwent RT using Superscript III according to manufacturer’s protocol using primer AR17 to amplify the Ag10 adapters and a primer to amplify GPx1. Following RT, qPCR was performed with a master mix (Thermo Fisher, 4385612) and using a Quantstudio 3 Real-Time PCR machine according to manufacturer specifications to quantify the accumulation of the NSD reporter normalized to GPx1 mRNA. All RT and qPCR primers are listed in Supplementary Table 2.

## Supplementary figures

**SUPPLEMENTARY FIGURE S1.**
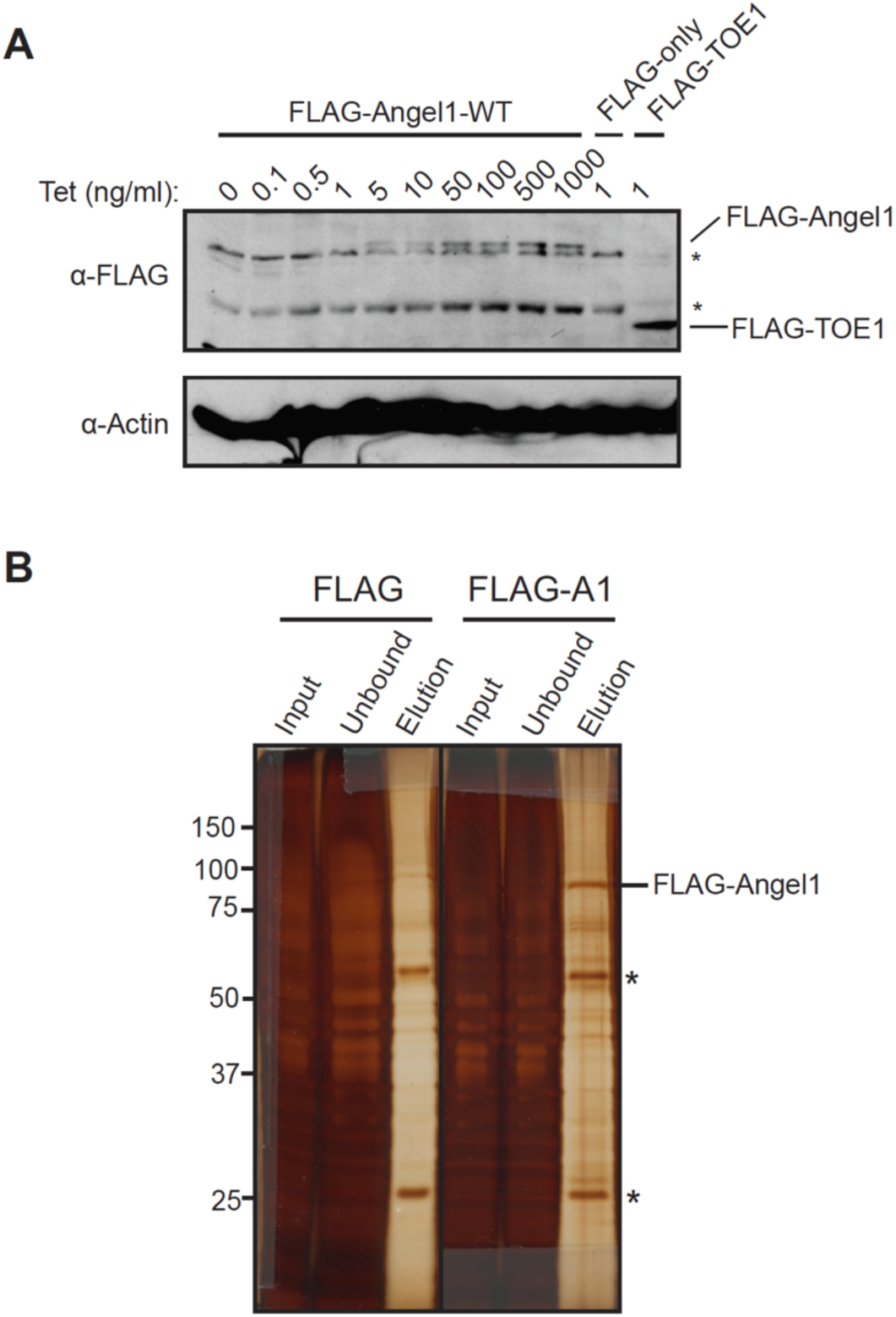
FLAG-Angel1 HEK293 Flp-In T-REx cell line validation. **(A)** Western blot showing expression of FLAG-Angel1 WT with titration of tetracycline (Tet) as indicated above lanes. The parental Flp-In T-REx line expressing no fusion protein serves as a negative control (FLAG-only), and Flp-In T-REx expressing FLAG-TOE1 at near endogenous levels was included as a reference. β-Actin was used as a loading control. *: non-specific bands. (B) Silver-stained SDS-PAGE gel examining protein from FLAG-Angel1-WT IPs. An input sample of 10% of the total cell extract used for the IP (Input), a sample from the remaining lysate after IP containing unbound proteins (Unbound), and a sample of the pooled elutions (Elution) were run. A control IP using a Flp-In T-REx line expressing no fusion protein (FLAG) was run alongside. *: denotes the antibody heavy and light chain.

**SUPPLEMENTARY FIGURE S2.**
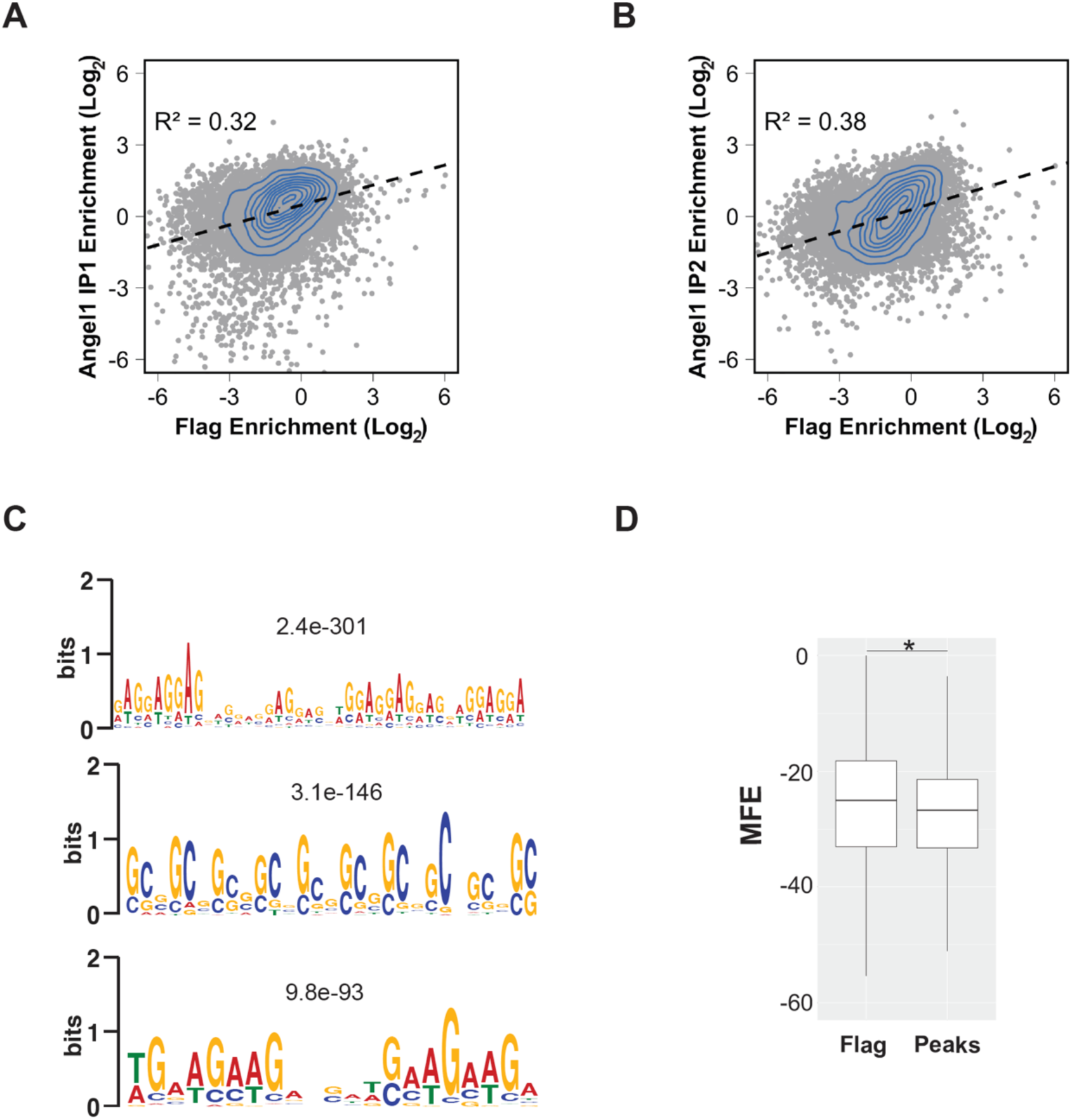
Extended eCLIP analysis. (A and B) Log_2_ fold enrichment of IP reads over input compared between either Flag-tagged Angel1 eCLIP replicate and the parental Flag control. Grey dots represent individual genes and the blue contour lines represent a 2D kernel density estimate. Pearson correlations are shown. (C) Logo plots of top three enriched nucleotide motifs identified by MEME analysis of sequences within 50 nucleotides upstream or downstream of Angel1 CLIP peaks as compared to sequences within 50 nucleotides of mapped reads in the FLAG input sample. E-values are displayed above each plot. (D) Box plots comparing predicted mean free energy (MFE) calculated by RNAfold for regions within 50 nucleotides upstream or downstream of Angel1 CLIP peaks and within 50 nucleotides of mapped reads in the FLAG input sample. Lower MFE scores are associated with stronger predicted secondary structure. *: p-value < 2.2e-22 (KS-test).

**SUPPLEMENTARY FIGURE S3.**
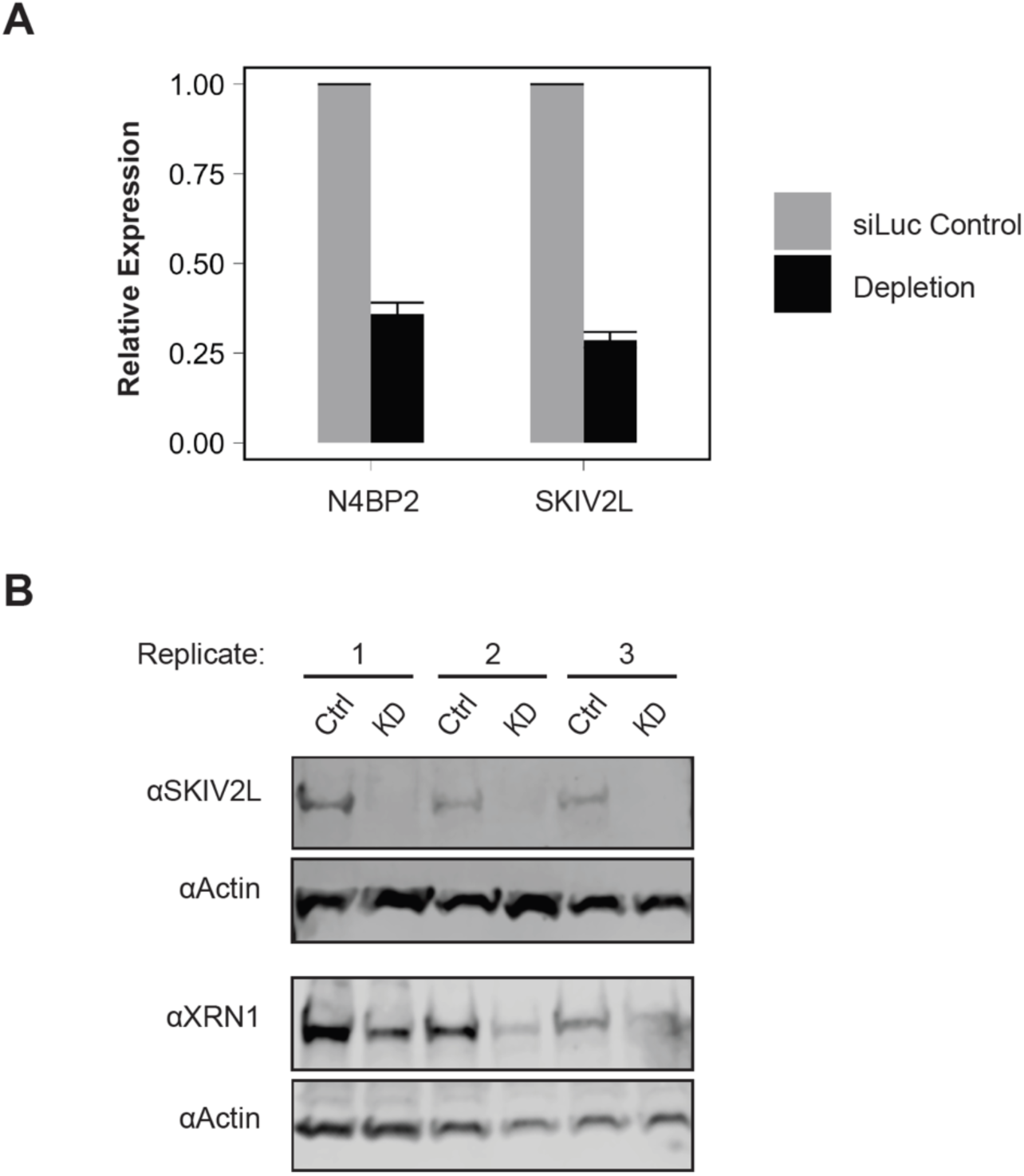
Validation of siRNA-mediated depletions. (A) Relative expression of knockdown targets compared to a control knockdown (siLuc) obtained from RT-qPCR. Error bars represent standard error of the mean (n=3). Comparisons between each knockdown and its control had a p-value < 0.05 as calculated by a one-tailed Student’s t-test. (B) Western blots examining levels of SKIV2L and XRN1 proteins after siRNA-mediated knockdown (KD) in triplicate or knockdown with a non-targeting control siRNA (Ctrl). β-Actin is shown as a loading control.

**SUPPLEMENTARY FIGURE S4.**
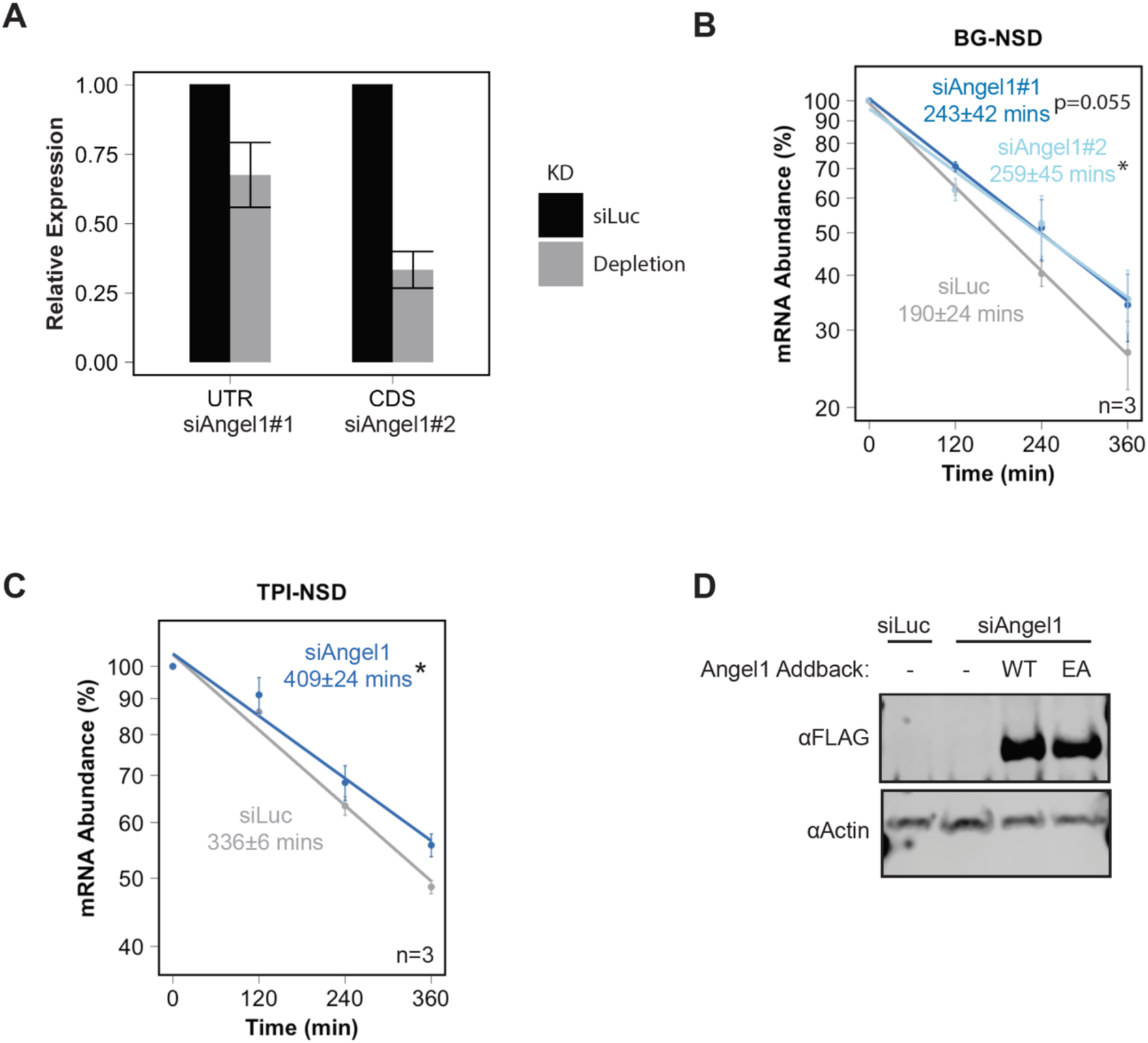
Additional BG-NSD validation. (A) Relative expression of Angel1 mRNA levels determined by RT-qPCR for two independent Angel1 siRNAs compared to a non-targeting control (siLuc). Comparisons between each knockdown and its control had a p-value of < 0.05 as calculated by a one-tailed Student’s t-test. Error bars represent standard error of the mean (n=3). siRNA#1, which targets endogenous but not exogenous Angel1, was used for all other depletions. (B) Exponential decay graphs of pulse-chase mRNA decay experiments using the BG-NSD substrate with depletion using the Angel1-targeting siRNAs shown in panel *A* and siLuc siRNA. (C) Exponential decay graphs of pulse-chase mRNA decay experiments using the TPI-NSD substrate with depletion of Angel1 or a non-targeting control. Error bars represent standard error of the mean. *: p-value < 0.05. n=3. (D) A Western blot examining total protein from Hela Tet-off cells either depleted by a non-targeting siRNA (siLuc) or an siRNA targeting endogenous Angel1 (siAngel1). Lanes 3 and 4 represent cells that were transfected with constructs expressing siRNA-resistant active (WT) or catalytically dead (EA) FLAG-Angel1 to complement depletion. β-Actin is shown as a loading control.

**SUPPLEMENTARY FIGURE S5.**
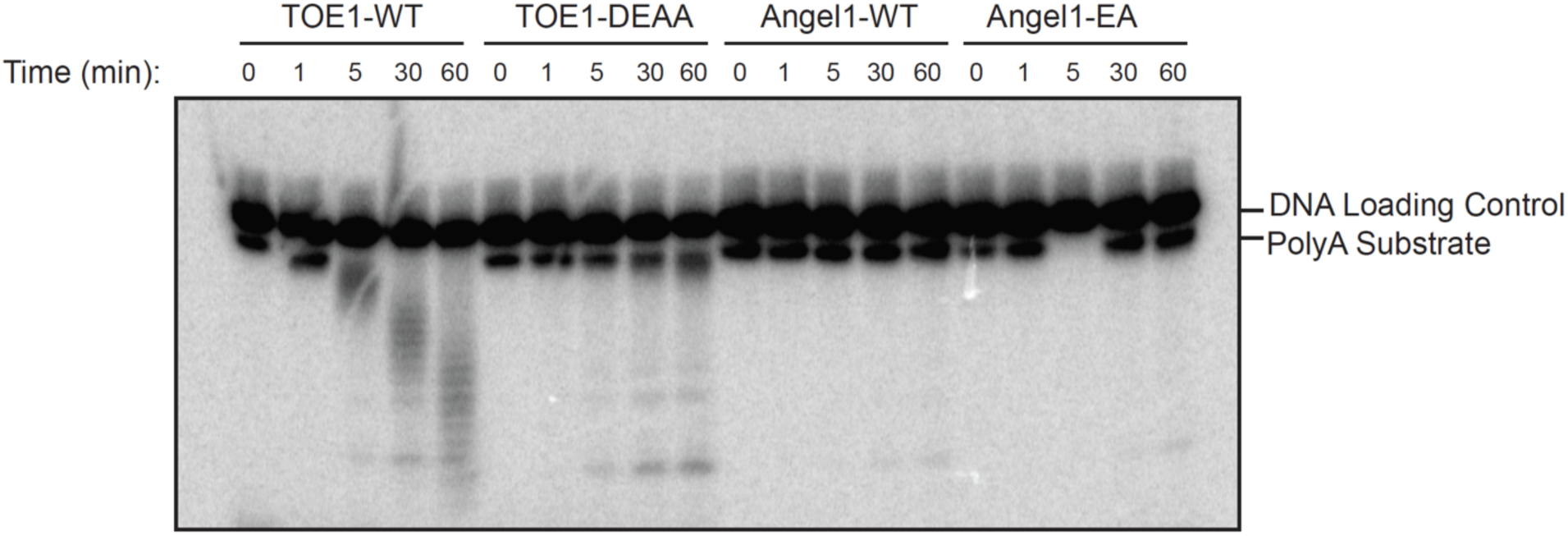
Angel1 shows no activity in a deadenylation assay. Phosphorimager scan of a 5’ ^32^P-labelled poly-A RNA substrate incubated with the indicated active (WT) or catalytically dead (DEAA, EA) proteins (≈50 nM) for the indicated amounts of time and subsequently separated in a denaturing gel. A 5’ ^32^P-labelled DNA substrate was included in each reaction as an internal loading control.

**SUPPLEMENTARY FIGURE S6.**
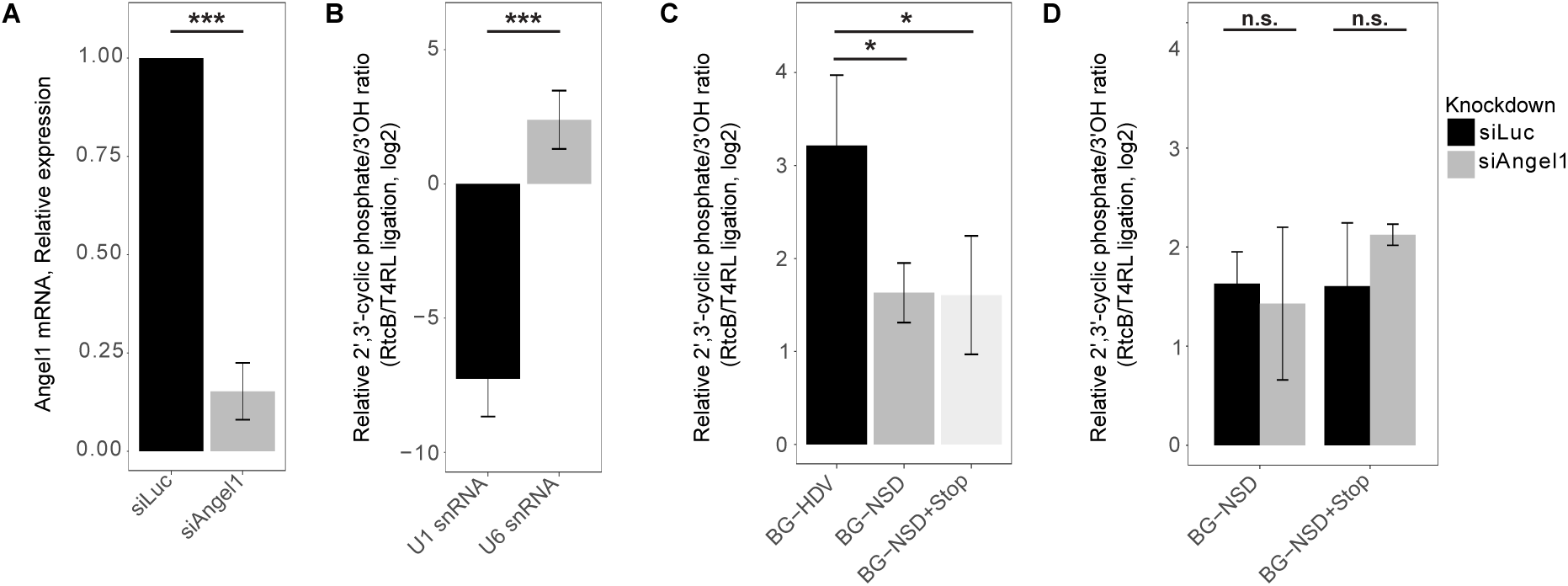
Angel1 depletion does not cause detectable accumulation of a 2’,3’ cyclic phosphate on the BG-NSD Reporter mRNA. (A) Relative levels of Angel1 mRNA determined by RT-qPCR of four independent samples after knockdown with siRNA for Angel1 (siAngel1#1) compared to non-targeting control siLuc. Error bars represent standard error of the mean (n=4). ***: p < 0.001, two-tailed Student’s t-test. (B) Relative 2’,3’ cyclic phosphate over 3’OH ratios of U1 (negative control) and U6 (positive control) snRNAs as measured by RNA adapter ligation by 2’,3’ cyclic phosphate-specific RtcB ligase as compared to 3’OH-specific T4 RNA ligase, followed by RT-qPCR. Error bars represent standard error of the mean (n=9). ***: p < 0.001, two-tailed Student’s t-test. (C) Relative 2’,3’ cyclic phosphate over 3’OH ratios of indicated β-globin reporter mRNAs measured as in panel *B*. Error bars represent standard error of the mean (n=3). *: p < 0.05, two-tailed Student’s t-test. (D) Relative 2’,3’ cyclic phosphate over 3’OH ratios of BG-NSD+Stop and BG-NSD reporter mRNAs in siLuc control depletion versus siAngel1 depletion conditions, measured as in panel *B* (n=3). n.s.: no significant difference by two-tailed Student’s t-test.

## References

Bailey TL, Boden M, Buske FA, Frith M, Grant CE, Clementi L, Ren J, Li WW, Noble WS. 2009. MEME SUITE: tools for motif discovery and searching. Nucleic Acids Research 37: W202– W208.

Bao C, Loerch S, Ling C, Korostelev AA, Grigorieff N, Ermolenko DN. 2020. mRNA stem-loops can pause the ribosome by hindering A-site tRNA binding. eLife 9: 1–67.

Bar-Peled L, Chantranupong L, Cherniack AD, Chen WW, Ottina KA, Grabiner BC, Spear ED, Carter SL, Meyerson M, Sabatini DM. 2013. A Tumor Suppressor Complex with GAP Activity for the Rag GTPases That Signal Amino Acid Sufficiency to mTORC1. Science 340: 1100–1106.

Cai W, Wei Y, Jarnik M, Reich J, Lilly MA. 2016. The GATOR2 Component Wdr24 Regulates TORC1 Activity and Lysosome Function ed. H. Kramer. PLoS Genetics 12: e1006036.

Chandrasekaran V, Juszkiewicz S, Choi J, Puglisi JD, Brown A, Shao S, Ramakrishnan V, Hegde RS. 2019. Mechanism of ribosome stalling during translation of a poly(A) tail. Nature Structural and Molecular Biology 26: 1132–1140.

Chen B, Retzlaff M, Roos T, Frydman J. 2011. Cellular strategies of protein quality control. Cold Spring Harbor Perspectives in Biology 3: 1–14.

Chyzyńska K, Labun K, Jones C, Grellscheid SN, Valen E. 2021. Deep conservation of ribosome stall sites across RNA processing genes. NAR Genomics and Bioinformatics 3: 1–13.

Clement SL, Lykke-Andersen J. 2008. A Tethering Approach to Study Proteins that Activate mRNA Turnover in Human Cells. Methods in Molecular Biology 419: 121–133.

Damgaard CK, Lykke-Andersen J. 2011. Translational coregulation of 5’TOP mRNAs by TIA-1 and TIAR. Genes and Development 25: 2057–2068.

Doma MK, Parker R. 2006. Endonucleolytic cleavage of eukaryotic mRNAs with stalls in translation elongation. Nature 440: 561–564.

Doma MK, Parker R. 2007. RNA Quality Control in Eukaryotes. Cell 131: 660–668.

D’Orazio KN, Green R. 2021. Ribosome states signal RNA quality control. Molecular Cell 81: 1372–1383.

D’Orazio KN, Wu CC-C, Sinha N, Loll-Krippleber R, Brown GW, Green R. 2019. The endonuclease Cue2 cleaves mRNAs at stalled ribosomes during No Go Decay. eLife 8.

Durand S, Franks TM, Lykke-Andersen J. 2016. Hyperphosphorylation amplifies UPF1 activity to resolve stalls in nonsense-mediated mRNA decay. Nature Communications 7: 1–12.

Durand S, Lykke-Andersen J. 2013. Nonsense-mediated mRNA decay occurs during eIF4F-dependent translation in human cells. Nature Structural and Molecular Biology 20: 702– 709.

Endoh T, Sugimoto N. 2016. Mechanical insights into ribosomal progression overcoming RNA G-quadruplex from periodical translation suppression in cells. Scientific Reports 6: 22719.

Ermolaeva MA, Dakhovnik A, Schumacher B. 2015. Quality control mechanisms in cellular and systemic DNA damage responses. Ageing research reviews 23: 3–11.

Franks TM, Singh G, Lykke-Andersen J. 2010. Upf1 ATPase-Dependent mRNP Disassembly Is Required for Completion of Nonsense-Mediated mRNA Decay. Cell 143: 938–950.

Frischmeyer PA, Van Hoof A, O’Donnell K, Guerrerio AL, Parker R, Dietz HC. 2002. An mRNA surveillance mechanism that eliminates transcripts lacking termination codons. Science 295: 2258–2261.

Garshott DM, An H, Sundaramoorthy E, Leonard M, Vicary A, Harper JW, Bennett EJ. 2021. iRQC, a surveillance pathway for 40S ribosomal quality control during mRNA translation initiation. Cell reports 36: 109642.

Glover ML, Burroughs AM, Monem PC, Egelhofer TA, Pule MN, Aravind L, Arribere JA. 2020. NONU-1 Encodes a Conserved Endonuclease Required for mRNA Translation Surveillance. Cell Reports 30: 4321–4331.e4.

Goldstrohm AC, Wickens M. 2008. Multifunctional deadenylase complexes diversify mRNA control. Nature reviews Molecular cell biology 9: 337–344.

Gosselin P, Martineau Y, Morales J, Czjzek M, Glippa V, Gauffeny I, Morin E, Le Corguillé G, Pyronnet S, Cormier P, et al. 2013. Tracking a refined eIF4E-binding motif reveals Angel1 as a new partner of eIF4E. Nucleic Acids Research 41: 7783–7792.

Gu J, Shumyatsky G, Makan N, Reddy R. 1997. Formation of 2’,3’-cyclic phosphates at the 3’ end of human U6 small nuclear RNA in vitro. Journal of Biological Chemistry 272: 21989– 21993.

Han P, Shichino Y, Schneider-Poetsch T, Mito M, Hashimoto S, Udagawa T, Kohno K, Yoshida M, Mishima Y, Inada T, et al. 2020. Genome-wide Survey of Ribosome Collision. Cell Reports 31: 107610.

Huter P, Arenz S, Bock L V., Graf M, Frister JO, Heuer A, Peil L, Starosta AL, Wohlgemuth I, Peske F, et al. 2017. Structural Basis for Polyproline-Mediated Ribosome Stalling and Rescue by the Translation Elongation Factor EF-P. Molecular Cell 68: 515–527.e6.

Ibrahim F, Maragkakis M, Alexiou P, Mourelatos Z. 2018. Ribothrypsis, a novel process of canonical mRNA decay, mediates ribosome-phased mRNA endonucleolysis. Nature Structural & Molecular Biology 25: 302–310.

Ikeuchi K, Tesina P, Matsuo Y, Sugiyama T, Cheng J, Saeki Y, Tanaka K, Becker T, Beckmann R, Inada T. 2019. Collided ribosomes form a unique structural interface to induce Hel2-driven quality control pathways. The EMBO Journal 38: e100276.

Joazeiro CAP. 2019. Mechanisms and functions of ribosome-associated protein quality control. Nature reviews Molecular cell biology 20: 368–383.

Jurkin J, Henkel T, Nielsen AF, Minnich M, Popow J, Kaufmann T, Heindl K, Hoffmann T, Busslinger M, Martinez J. 2014. The mammalian tRNA ligase complex mediates splicing of XBP1 mRNA and controls antibody secretion in plasma cells. The EMBO Journal 33: 2922.

Juszkiewicz S, Chandrasekaran V, Lin Z, Kraatz S, Ramakrishnan V, Hegde RS. 2018. ZNF598 Is a Quality Control Sensor of Collided Ribosomes. Molecular cell 72: 469–481.

Kalisiak K, Kuliński TM, Tomecki R, Cysewski D, Pietras Z, Chlebowski A, Kowalska K, Dziembowski A. 2017. A short splicing isoform of HBS1L links the cytoplasmic exosome and SKI complexes in humans. Nucleic acids research 45: 2068–2080.

Kikin O, D’Antonio L, Bagga PS. 2006. QGRS Mapper: A web-based server for predicting G-quadruplexes in nucleotide sequences. Nucleic Acids Research 34: W676–W682.

Kowalsky AH, Namkoong S, Mettetal E, Park HW, Kazyken D, Fingar DC, Lee JH. 2020. The GATOR2–mTORC2 axis mediates Sestrin2-induced AKT Ser/Thr kinase activation. The Journal of Biological Chemistry 295: 1769.

Kurzik-Dumke U, Zengerle A. 1996. Identification of a novel Drosophila melanogaster gene, angel, a member of a nested gene cluster at locus 59F4,5. Biochimica et Biophysica Acta 1308: 177–181.

Lardelli RM, Schaffer AE, C Eggens VR, Zaki MS, Grainger S, Sathe S, Van Nostrand EL, Schlachetzki Z, Rosti B, Akizu N, et al. 2017. Biallelic mutations in the 3ʹ exonuclease TOE1 cause pontocerebellar hypoplasia and uncover a role in snRNA processing. Nature Genetics 49: 457–464.

Liao Y, Smyth GK, Shi W. 2014. featureCounts: an efficient general purpose program for assigning sequence reads to genomic features. Bioinformatics 30: 923–930.

Lykke-Andersen J, Bennett EJ. 2014. Protecting the proteome: Eukaryotic cotranslational quality control pathways. Journal of Cell Biology.

Lykke-Andersen J, Shu M-D, Steitz JA. 2000. Human Upf Proteins Target an mRNA for Nonsense-Mediated Decay When Bound Downstream of a Termination Codon. Cell 103: 1121– 1131.

Nagaraj N, Wisniewski JR, Geiger T, Cox J, Kircher M, Kelso J, Pääbo S, Mann M. 2011. Deep proteome and transcriptome mapping of a human cancer cell line. Molecular systems biology 7.

Navickas A, Chamois S, Saint-Fort R, Henri J, Torchet C, Benard L. 2020. No-Go Decay mRNA cleavage in the ribosome exit tunnel produces 5ʹ-OH ends phosphorylated by Trl1. Nature Communications 11: 122.

Pinto PH, Kroupova A, Schleiffer A, Mechtler K, Jinek M, Weitzer S, Martinez J. 2020. ANGEL2 is a member of the CCR4 family of deadenylases with 2’,3’-cyclic phosphatase activity. Science (New York, NY) 369: 524–530.

Pisareva VP, Skabkin MA, Hellen CUT, Pestova T V, Pisarev A V. 2011. Dissociation by Pelota, Hbs1 and ABCE1 of mammalian vacant 80S ribosomes and stalled elongation complexes. The EMBO journal 30: 1804–17.

Reinke AW, Mak R, Troemel ER, Ben EJ. 2017. In vivo mapping of tissue- and subcellular-specific proteomes in Caenorhabditis elegans. Science Advances 3.

Rodriguez JM, Maietta P, Ezkurdia I, Pietrelli A, Wesselink JJ, Lopez G, Valencia A, Tress ML. 2013. APPRIS: annotation of principal and alternative splice isoforms. Nucleic Acids Research 41: D110.

Saito S, Hosoda N, Hoshino SI. 2013. The Hbs1-Dom34 protein complex functions in non-stop mRNA decay in mammalian cells. Journal of Biological Chemistry 288: 17832–17843.

Shigematsu M, Kirino Y. 2020. Oxidative stress enhances the expression of 2ʹ,3ʹ-cyclic phosphate-containing RNAs. RNA Biology 17: 1060–1069.

Simms CL, Hudson BH, Mosior JW, Rangwala AS, Zaher HS. 2014. An active role for the ribosome in determining the fate of oxidized mRNA. Cell reports 9: 1256–64.

Simms CL, Yan LL, Zaher HS. 2017. Ribosome Collision Is Critical for Quality Control during No-Go Decay. Molecular Cell 68: 361–373.e5.

Singh G, Jakob S, Kleedehn MG, Lykke-Andersen J. 2007. Communication with the Exon-Junction Complex and Activation of Nonsense-Mediated Decay by Human Upf Proteins Occur in the Cytoplasm. Molecular Cell 27: 780–792.

Singh G, Rebbapragada I, Lykke-Andersen J. 2008. A competition between stimulators and antagonists of Upf complex recruitment governs human nonsense-mediated mRNA decay. PLoS biology 6: 860–871.

Sundaramoorthy E, Leonard M, Mak R, Liao J, Fulzele A, Bennett EJ. 2017. ZNF598 and RACK1 Regulate Mammalian Ribosome-Associated Quality Control Function by Mediating Regulatory 40S Ribosomal Ubiquitylation. Molecular Cell 65: 751–760.e4.

Sundaramoorthy E, Ryan AP, Fulzele A, Leonard M, Daugherty MD, Bennett EJ. 2021. Ribosome quality control activity potentiates vaccinia virus protein synthesis during infection. Journal of cell science 134.

Tropea D, Hardingham N, Millar K, Fox K. 2018. Mechanisms underlying the role of DISC1 in synaptic plasticity. The Journal of physiology 596: 2747–2771.

Tsuboi T, Kuroha K, Kudo K, Makino S, Inoue E, Kashima I, Inada T. 2012. Dom34:Hbs1 Plays a General Role in Quality-Control Systems by Dissociation of a Stalled Ribosome at the 3ʹ End of Aberrant mRNA. Molecular Cell 46: 518–529.

Tuck AC, Rankova A, Arpat AB, Liechti LA, Hess D, Iesmantavicius V, Castelo-Szekely V, Gatfield D, Bühler M. 2020. Mammalian RNA Decay Pathways Are Highly Specialized and Widely Linked to Translation. Molecular Cell 77: 1222–1236.e13.

Van Hoof A, Frischmeyer PA, Dietz HC, Parker R. 2002. Exosome-mediated recognition and degradation of mRNAs lacking a termination codon. Science 295: 2262–2264.

Van Nostrand EL, Nguyen TB, Gelboin-Burkhart C, Wang R, Blue SM, Pratt GA, Louie AL, Yeo GW. 2017. Robust, Cost-Effective Profiling of RNA Binding Protein Targets with Single-end Enhanced Crosslinking and Immunoprecipitation (seCLIP). pp. 177–200, Humana Press, New York, NY.

Van Nostrand EL, Pratt GA, Shishkin AA, Gelboin-Burkhart C, Fang MY, Sundararaman B, Blue SM, Nguyen TB, Surka C, Elkins K, et al. 2016. Robust transcriptome-wide discovery of RNA-binding protein binding sites with enhanced CLIP (eCLIP). Nature Methods 13: 508– 514.

Wagner E, Clement SL, Lykke-Andersen J. 2007. An unconventional human Ccr4-Caf1 deadenylase complex in nuclear cajal bodies. Molecular and cellular biology 27: 1686– 1695.

Wang H, Morita M, Yang X, Suzuki T, Yang W, Wang J, Ito K, Wang Q, Zhao C, Bartlam M, et al. 2010. Crystal structure of the human CNOT6L nuclease domain reveals strict poly(A) substrate specificity. The EMBO journal 29: 2566–2576.

Weber R, Chung MY, Keskeny C, Zinnall U, Landthaler M, Valkov E, Izaurralde E, Igreja C. 2020. 4EHP and GIGYF1/2 Mediate Translation-Coupled Messenger RNA Decay. Cell Reports 33: 108262.

Yang W. 2011. Nucleases: Diversity of structure, function and mechanism. Quarterly Reviews of Biophysics 44: 1–93.

Yi H, Park J, Ha M, Lim J, Chang H, Kim VN. 2018. PABP Cooperates with the CCR4-NOT Complex to Promote mRNA Deadenylation and Block Precocious Decay. Mol Cell 70: 1081–1088.e5.

Yip MCJ, Keszei AFA, Feng Q, Chu V, McKenna MJ, Shao S. 2019. Mechanism for recycling tRNAs on stalled ribosomes. Nature Structural & Molecular Biology 26: 343–349.

Zillmann M, Gorovsky MA, Phizicky EM. 1991. Conserved mechanism of tRNA splicing in eukaryotes. Molecular and Cellular Biology 11: 5410.

